# The CLIC/GEEC pathway regulates particle endocytosis and formation of the virus-containing compartment (VCC) in HIV-1-infected macrophages

**DOI:** 10.1101/2024.09.05.611375

**Authors:** Kathleen Candor, Lingmei Ding, Sai Balchand, Jason E. Hammonds, Paul Spearman

**Affiliations:** Immunology Graduate Program, University of Cincinnati, and Infectious Diseases Division, Cincinnati Children’s Hospital, Cincinnati, OH USA; Infectious Diseases Division, Cincinnati Children’s Hospital and University of Cincinnati, Cincinnati, OH USA

**Author notes:** Corresponding author: Infectious Diseases, Cincinnati Children’s Hospital, 3333 Burnet Avenue, MLC 7017, Cincinnati, OH USA 45229.

**Keywords:** HIV-1, macrophage, virus-containing compartment, CLIC/GEEC, dynamin, endocytosis, Siglec-1, CD98

## Abstract

HIV-1 particles are captured by the immunoglobulin superfamily member Siglec-1 on the surface of macrophages and dendritic cells, leading to particle internalization and facilitating trans-infection of CD4+ T cells. HIV-1-infected macrophages develop a unique intracellular compartment termed the virus-containing compartment (VCC) that exhibits characteristic markers of the late endosome and is enriched in components of the plasma membrane (PM). The VCC has been proposed as the major site of particle assembly in macrophages. Depleting Siglec-1 from macrophages significantly reduces VCC formation, implying a link between the capture and endocytosis of external HIV-1 particles and the development of VCCs within HIV-infected cells. We found that endocytosis of particles to the VCC was independent of clathrin, but required dynamin-2. CD98 and CD44, classical markers of the CLIC/GEEC pathway, colocalized with Siglec-1 and HIV-1 particles within the VCC. Inhibition of CLIC/GEEC-mediated endocytosis by chemical or genetic means resulted in the arrest of captured HIV-1 particles on the macrophage cell surface and prevented VCC formation. Virus-like particles (VLPs) were taken up within CD98 and Siglec-1-enriched tubular membranes that migrated centripetally over time to form VCC-like structures. These findings indicate that following capture of virus by Siglec-1, particles follow an endocytic route to the VCC that requires both the CLIC/GEEC pathway and dynamin-2. We propose a model in which internalization of HIV-1 particles together with CLIC/GEEC membranes leads to the formation of the VCC in HIV-infected macrophages, creating an intracellular platform that facilitates further particle assembly and budding.

**Author Summary:** The major cell types infected by HIV are CD4+ T cells and macrophages. Infection of macrophages is of great interest because this cell type can contribute to transmission of virus within tissues, and because infected macrophages contribute to HIV-related complications that include neurologic disorders, endocrine disorders, and cardiovascular disease. HIV infection of macrophages may also create a latent reservoir that persists in infected individuals despite administration of antiretroviral therapy. Here we focused on a compartment that forms in HIV-infected macrophages termed the virus-containing compartment or VCC. The VCC is an intracellular compartment that has been described as a site of assembly and as a holding compartment for viruses, and contains components of both intracellular organelles and the plasma membrane. We identified an endocytosis pathway that helps to explain the origins of the VCC, the CLIC/GEEC pathway. Inhibition of the CLIC/GEEC pathway prevented virus-like particle delivery to the VCC and prevented the VCC from forming in HIV-infected macrophages. The GTPase dynamin-2 was also required for delivery of HIV particles to the VCC. This study identifies a new facet of how HIV interacts with macrophages, suggesting that disruption of this pathway could be a therapeutic strategy with implications for HIV cure efforts.

## Introduction

Tissue macrophages are an important target of HIV-1 infection. Infected macrophages have been found in numerous tissues derived from HIV-infected individuals, including peripheral lymph nodes, gut-associated lymphoid tissue, liver, lung, genitourinary tract, bone, and the brain (1–8). Infected macrophages are thought to play a prominent role in HIV-related comorbidities, including HIV-associated neurocognitive disorders (HAND), accelerated cardiovascular disease, early immune aging, and endocrine disorders (9–13). Infected macrophages may also form a long-lived reservoir for HIV-1 (14–16). It is therefore important to understand how HIV-1 interacts with and alters macrophage structure and function.

HIV-1-infected macrophages form an unique intracellular structure known as the virus-containing compartment (VCC), also referred to as the intracellular plasma membrane compartment (IPMC) (17–20). The VCC shares many molecular features of the late endosome/multivesicular body (MVB), including the presence of MHC class II, LAMP-1, and tetraspanins CD9, CD37, CD53, CD63, CD81, and CD82. The VCC is enriched in β2 integrins, the scavenger receptor CD36, hyaluronate receptor/surface glycoprotein CD44, and phosphatidylinositol 4,5 bisphosphate (PIP_2_) (17, 21–23). In contrast to the late endosome, the VCC has a near neutral pH (18). Tubular connections extending from the VCC to the extracellular milieu have been observed by fluorescence microscopy, transmission EM, and ion abrasion scanning EM (17, 20, 23, 24). The origin and function of these tubules of the VCC are unknown, although it has been suggested they could represent an exit route for viruses from the VCC (24). The VCC may serve as a virion storage site for HIV-1 that can promote trans-infection upon contact with target T cells or uninfected macrophages, similar to the role a similar “holding” compartment plays in dendritic cells (DCs) (25, 26). The presence of budding forms observed by EM on VCC membranes, and the general paucity of particles seen assembling on the PM of macrophages, has led to the conclusion that the VCC is an intracellular assembly compartment and the major site of assembly in macrophages (17, 18, 22, 27). However, the genesis of an intracellular membranous compartment made up of both late endosomal and plasma membrane components remains uncertain.

Siglec-1 was identified as an important myeloid cell lectin that binds and captures HIV-1 particles through interactions with gangliosides on the lipid envelope of the virus, contributing to trans-infection of target cells by DCs (28, 29). We have previously shown that Siglec-1 on the surface of monocyte-derived macrophages (MDMs) captures exogenous non-infectious HIV-1 virus-like particles (VLPs) and leads to their internalization into the macrophage to form a structure identical to the VCC in morphology, location, and in the presence of typical VCC markers (30). Furthermore, when HIV-1 VLPs are added exogenously to infected cells that contain a pre-formed VCC, they are delivered in a Siglec-1-dependent manner into the authentic VCC. Siglec-1 depletion inhibits VCC formation within infected MDMs and prevents a VCC-like structure from forming following exogenous addition of VLPs, suggesting a common role for Siglec-1 in VCC formation. Capture of HIV-1 on the surface of activated DCs leads to nanoclustering of Siglec-1, and this nanoclustering together with activation of the actin cytoskeleton is required for internalization to the VCC (31). However, it remains unknown how the development of virus-Siglec-1 nanoclusters leads to particle uptake or endocytosis of particles to the VCC.

Here we evaluated the role of cellular endocytic pathways in the uptake of HIV-1 particles following Siglec-1-mediated particle capture. Inhibition of dynamin-2 through chemical and genetic means resulted in the arrest of captured particles and Siglec-1 in a more peripheral location in MDMs, preventing VCC formation. Inhibition of clathrin-mediated endocytosis did not arrest VCC formation, leading us to focus on clathrin-independent pathways. Remarkably, clathrin-independent carriers/glycosylphosphatidylinositol-anchored protein-enriched endocytic compartments (CLIC/GEEC) cargo were highly enriched in the VCC of infected cells, and disruption of the CLIC/GEEC pathway prevented VLP uptake in infected cells and disrupted VCC formation in infected MDMs. Furthermore, tubular membranes enriched in CLIC/GEEC cargo protein CD98 were found to contain Siglec-1-captured VLPs. Migration of Siglec-1, CLIC/GEEC cargo proteins, and VLPs to the center of the cell over time resulted in the formation of structures identical to the authentic VCCs of HIV-1-infected macrophages. These results demonstrate that the CLIC/GEEC pathway is involved in the movement of Siglec-1-capture particles to the VCC and is required for VCC formation in infected MDMs.

## Results

### Dynamin inhibition or depletion prevents VLP uptake and VCC formation in MDMs

We first asked if dynamin is required for VLP uptake and VCC formation following particle capture by Siglec-1. The dynasore analog, Dyngo4a, has been shown to exhibit specific and potent inhibition of dynamin-mediated endocytosis (32, 33). We therefore examined the effects of Dyngo4a treatment on HIV-1 VLP uptake into a perinuclear VCC in uninfected MDMs. MDMs were incubated with GFP-tagged HIV-1 VLPs for 5 hours to allow a VCC-like structure to form, after which they were treated with Dyngo4a or DMSO control for 30 minutes. mCherry-tagged VLPs were then added to the MDMs to assess the effects of dynamin inhibition on VLP uptake into the VCC. Transferrin was added 30 minutes prior to fixation to monitor dynamin-dependent uptake. As expected, Dyngo4a-treated cells were deficient in uptake of transferrin as compared with control cells (Figure 1A, 1B, transferrin panels, with quantitation in 1D). HIV-1 mcherry VLP uptake was reduced but not eliminated by Dyngo4a treatment (Figure 1A, 1B, and volume measurement in 1C). The VLPs that did enter the cell largely failed to reach the VCC following Dyngo4a treatment (Figure 1A, 1B, and colocalization data in 1E), while in control cells mcherry VLPs colocalized extensively with GPF VLPs within the VCC. The HIV-1 mcherry VLPs in Dyngo4a-treated cells appeared to enter the cell, but were located in a position superior to the VCC as if arrested in transit. Notably, Dyngo4a treatment specifically blocked dynamin-dependent endocytosis in MDMs, while not showing off-target effects on clathrin-independent endocytosis pathways, such as CLIC/GEEC and macropinocytosis, as indicated by preserved uptake of CD98 and low molecular weight dextran (Supplemental Figure 1A-1D). These results suggest that HIV-1 virion uptake into the VCC is dynamin-dependent, while in the absence of dynamin activity VLPs are still captured, but are aborted in reaching the VCC.

**Figure 1.**
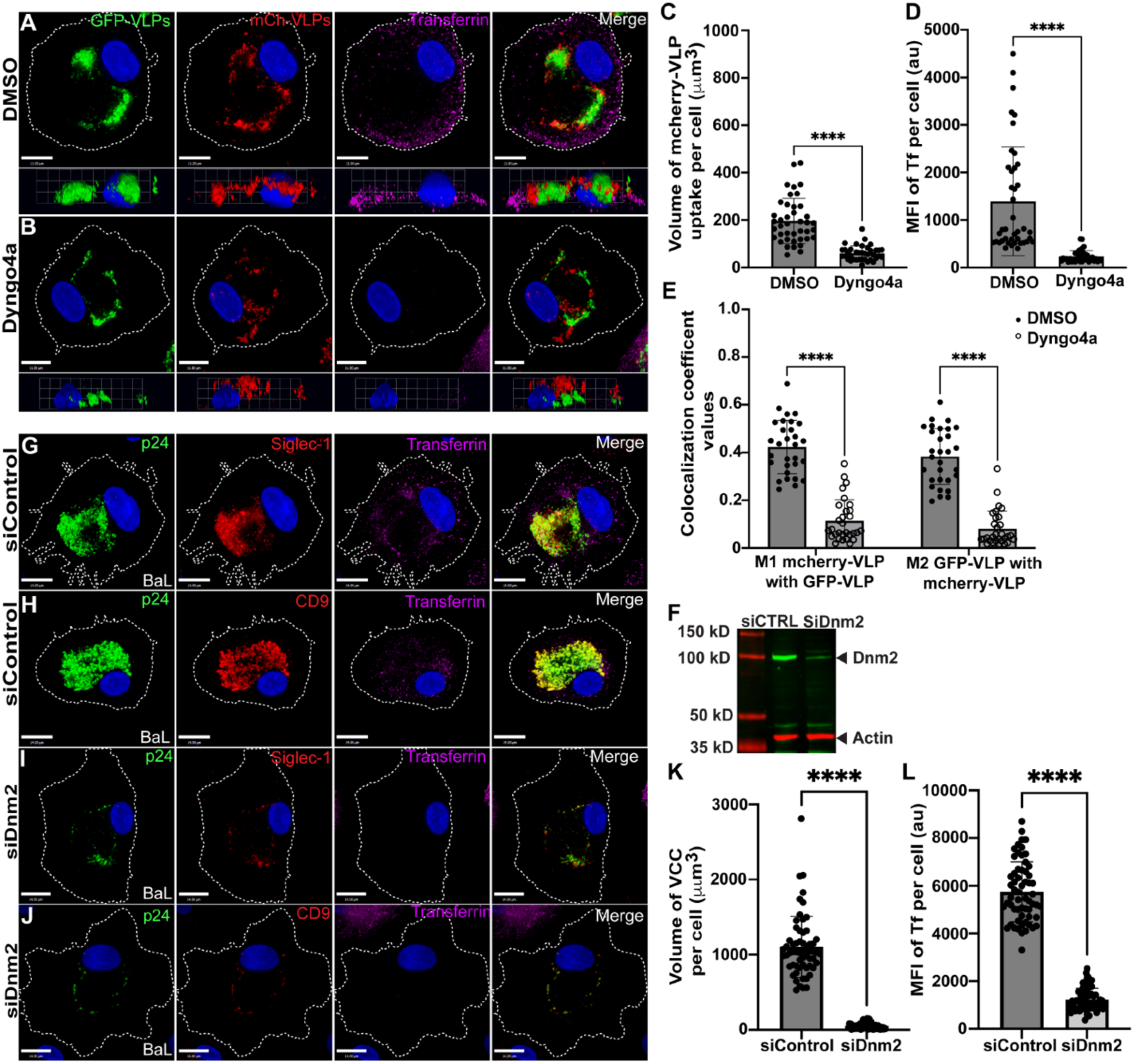
Dynamin plays an important role in VLP uptake and VCC formation in MDMs. (**A**, and **B**) Representative maximal intensity projections derived from z stacks of the MDMs, along with 3D side projection projection of the same images. (**A** and **B**) HIV-1 Gag-GFP VLPs were added to MDM cultures; following a 5 hour incubation they were treated with either DMSO (**A)** or Dyngo4a (**B**), followed by addition of HIV-1 Gag-mCherry VLPs for 2.5 hrs (red). Transferrin uptake (magenta) was evaluated during the final 30 minutes before fixation. Size bar = 11 μm. (**C**) Mean ± SD volume of mcherry-VLP uptake per cell (μ𝑚^!^) from a total of 30 MDMs in each group. (**D**) Mean ± SD fluorescence intensity of transferrin uptake per cell. (**E**) Colocalization coefficient values for mcherry VLP/ GFP VLP (M1) and GFP VLP/mcherry VLP (M2). (**F**) Representative knockdown efficiency of Dnm2 in MDMs with either siControl or siDnm2. (**G** and **H**) BaL-infected MDMs transfected with 50nM control siRNA. (**I** and **J**) BaL-infected MDMs transfected with 50nM dynamin-2 siRNA. Transferrin uptake is shown in magenta, p24 staining in green, and Siglec-1 or CD9 in red. Scale bar = 14 μm. (**K**) Mean ± SD volume of the VCC per BaL-infected cell (60 different BaL-infected MDMs per group). (**L**) Mean ± SD fluorescence intensity of transferrin uptake per infected cell. Data are representative of least three independent experiments. ****p <0.001 and ns, not significant (student’s t-test).

To further evaluate the role of dynamin in particle uptake and VCC formation, we utilized siRNA-mediated depletion of dynamin-2. We focused on dynamin 2 as it is ubiquitously expressed in all cells, whereas dynamin-1 is expressed in primarily in neurons and dynamin-3 is expressed primarily in brain, lung and testis (34, 35). MDMs were infected with HIV-1_BAL_ and treated with control siRNA or dynamin-2 siRNA, and VCC formation evaluated on day 12 post-infection. Dynamin-2 expression levels were reduced by greater than 70 percent with dynamin-2 siRNA (Figure 1F). The effectiveness of dynamin inhibition was documented through addition of transferrin on day 12 post-infection, as uptake was inhibited in dynamin-2 siRNA cells as compared with controls (Figure 1G-1J, with quantitation shown in 1L). Control siRNA-treated MDMs formed deep VCCs, as indicated by colocalization of HIV-1 particles, Siglec-1, and CD9 in a deep location in the cell (Figure 1G and 1H). Dynamin-2 depletion resulted in significant reduction in the VCC volume within HIV-1 infected MDMs (Figure 1I and 1J, with quantitation in 1K). The mean VCC volume from BaL infection was reduced from 1104 ± 52 mm^!^ to a mean of 53.55 ± 5 mm^!^ (Figure 1K). Together, chemical inhibition experiments and dynamin-2 depletion experiments support an important role for dynamin-2 in endocytosis of Siglec-1-captured particles and in VCC formation.

### Clathrin-mediated endocytosis (CME) is not required for Siglec-1-mediated particle uptake and VCC formation in macrophages

Endocytosis through CME requires dynamin for the scission and release of clathrin-coated vesicles from the plasma membrane (36, 37). To evaluate the potential role of CME in VLP uptake, we initially performed chemical inhibition experiments using Pitstop 2.0. However, this inhibitor demonstrated significant off-target effects in MDMs, including disruption of clathrin-independent endocytosis (CIE) pathways (Supplemental Figure 1E and 1F) as been previously published, limiting its utility for our studies (38). Therefore, we relied on depletion of key components of the CME pathway, epidermal growth factor receptor substrate 15 (Eps15) and Fer/CIPa homology domain protein 2 (FCHO2), both of which contribute to early steps of the CME pathway (36, 37, 39). Eps15 and FCHO2 expression levels in MDMs were reduced by more than 70 percent following siRNA-mediated depletion (Figure 2A and 2B). On day 6 following siRNA treatment, HIV-1 VLPs were added and allowed to be endocytosed for 15 hours. Transferrin was added 30 minutes prior to fixation to monitor CME-mediated uptake. Control siRNA-treated MDMs demonstrated robust uptake of transferrin (Figure 2C, with quantitation in 2G). As expected, depletion of either Eps15 and FCHO2 resulted in a significant reduction of transferrin uptake (Figure 2D and 2E, with quantitation in 2G). Despite robust inhibition of CME, there was no significant difference in the volume of the VLPs uptake in MDMs depleted of Eps15 or FCHO2 vs. control MDMs (Figure 2C, 2D and 2E, with quantitation in 2F). To further assess the role of the CME pathway in VCC formation, we performed depletion of Eps15 and FCHO2 in HIV-1-infected MDMs. We first infected the MDMs overnight with VSV-G-pseudotyped *vpu*-deficient NLU_del_, then utilized siRNA against Eps15, FCHO2 or control siRNA as before. Control siRNA-treated MDMs formed VCCs deep within the cell body and demonstrated robust transferrin uptake (Figure 2H, with quantitation in 2L). Depletion of Eps15 or FCHO2 led to inhibition of transferrin uptake as expected (Figure 2I and 2J and quantitation shown in 2L). No significant change in VCC volume was seen following depletion of either component of the CME pathway (Figure 2I and 2J, quantitation in 2K). These data suggest that the CME pathway is not involved in the formation of the VCC in HIV-infected MDMs. Based on these results, we turned our attention to clathrin-independent pathways of endocytosis.

**Figure 2.**
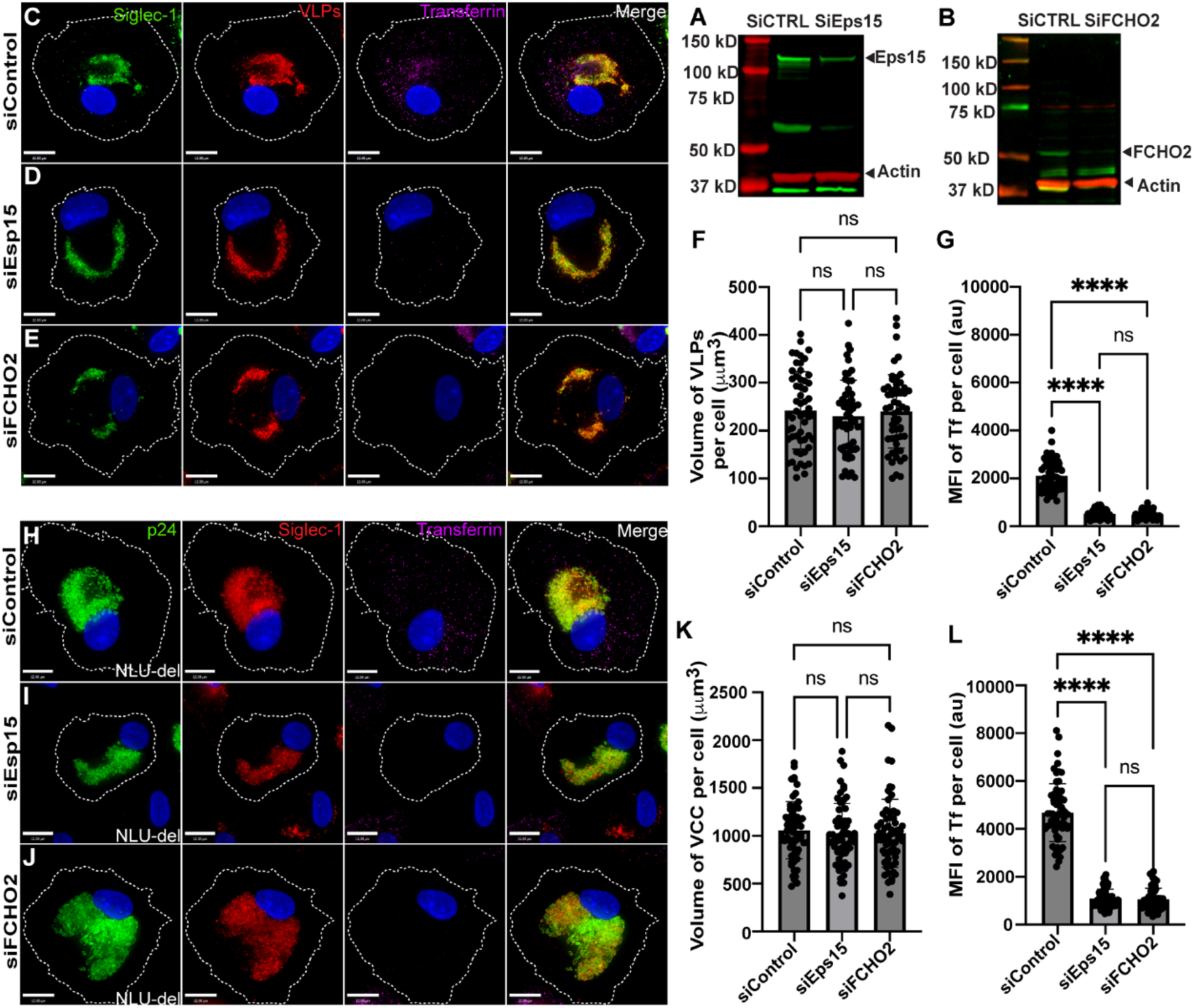
Inhibition of CME does not prevent formation of the VCC in MDMs. (**A**) Representative knockdown of Eps15 in MDMs transfected with siControl or siEps15. (**B**) Representative knockdown of FCHO2 in MDMs transfected with siControl or siFCHO2. (**C**) MDMs transfected with 50nM of control siRNA, showing uptake of mcherry VLPs (red) and transferrin (magenta) together with Siglec-1 (green). (**D**) MDMs transfected with siEps15 and stained as described for (C). **(E)** MDMs transfected with siFCHO2 and stained as described for (C). (**F**) Mean ± SD volume of VLPs per cell from 56 different MDMs for each experimental group. (**G**) Mean ± SD fluorescence intensity of transferrin per cell. **(H)** NL_Udel_-infected MDMs transfected with control siRNA, showing p24 (green), Siglec-1 (red), and transferrin (magenta). Scale bar = 12 μm. (**I)** NL_Udel_-infected MDMs transfected with Eps15 siRNA and stained as before. (**J**) NL_Udel_-infected MDMs transfected with FCHO2 siRNA. Scale bar =12 μm. (**K**) Mean ± SD volume of VCC per cell from a total of 60 different NL_Udel_-infected MDMs in each group. (**L**) Mean ± SD fluorescence intensity of transferrin uptake per cell. Data are representative of a minimum of three independent experiments ****p <0.001 and ns, not significant (Kruskal-Wallis test).

### The VCC in HIV-1 infected human MDMs is enriched in CLIC/GEEC cargo CD44 and CD98

After finding that clathrin-mediated endocytosis was not required for VCC formation, we next turned to an examination of clathrin-independent pathways. The hyaluronate receptor and surface glycoprotein CD44 has previously been described as a component of the VCC in infected MDMs (23). Because CD44 is a characteristic cargo protein of the CLIC/GEEC endocytosis pathway (40, 41), we hypothesized that the CLIC/GEEC pathway may play a role in VCC formation. We examined the subcellular distribution of another characteristic CLIC/GEEC cargo, the amino acid transporter CD98. MDMs were infected with HIV-1_BAL_ and incubated for 10 days to allow VCC formation. Established markers of the VCC, CD9 and Siglec-1, colocalized with virus in the VCC as expected (Figure 3A and 3B). CD44 was highly concentrated in the VCC as previously described (Figure 3C). Notably, CD98 was also highly enriched in the VCC, identifying a second classic marker of the CLIC/GEEC pathway in this compartment (Figure 3D). These findings showing enrichment of CLIC/GEEC cargo in the VCC then led us to further examine the potential functional role of this pathway in the formation of the VCC.

**Figure 3.**
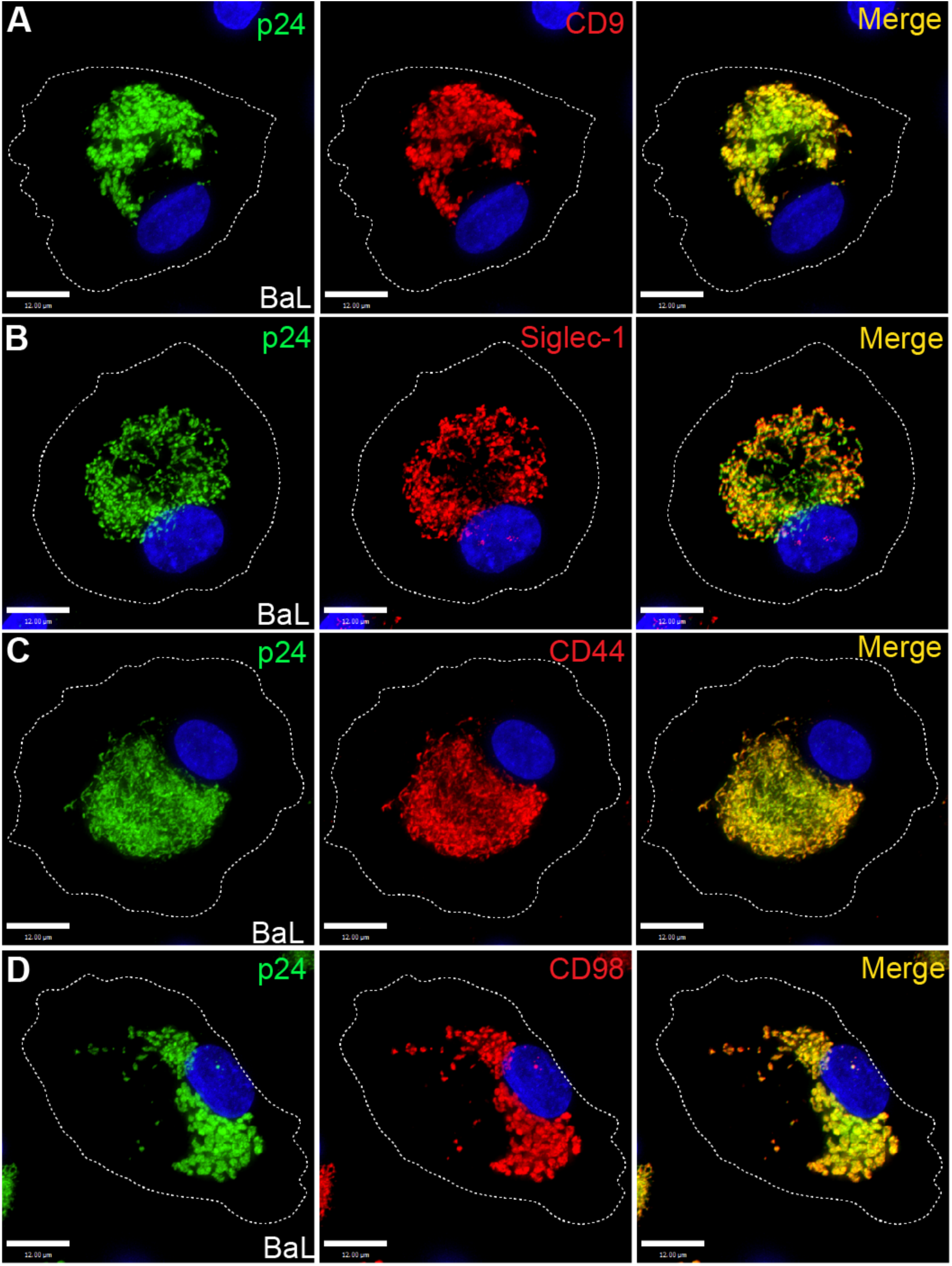
CLIC/GEEC Cargo in the VCC in HIV-1 BaL-infected MDMs. On day 12 post-infection, MDMs were fixed and immunostained for p24 (green) and **(A)** CD9, **(B)** Siglec-1, **(C)** CD44, or **(D)** CD98 (red). Scale bar = 12 μm.

### Chemical inhibition of the CLIC/GEEC pathway inhibits VLP uptake into MDMs

We next examined the effect of chemical inhibition of the CLIC/GEEC pathway using 7-keto-cholesterol (7-KC). 7-KC is an oxysterol that prevents close packing of acyl chains and has been shown to inhibit endocytosis via the CLIC/GEEC pathway (42, 43). MDMs were pre-treated with 7-KC prior to the addition of fluorescent HIV-1 VLPs, and endocytosis of VLPs compared with that seen in control cells. To monitor CLIC/GEEC inhibition, we utilized a pulse-chase with anti-CD44 or anti-CD98 antibodies, applied 30 minutes prior to fixation. Control MDMs readily took up VLPs into VCCs, where they colocalized with Siglec-1, CD98, and CD44 (Figure 4A and 4B, quantitation in 4E-4I). Treatment with 7-KC reduced the amount of CD98 antibody and CD44 antibody internalized, as expected, and also markedly inhibited VLP uptake. After treatment with 7-KC, VLPs captured by Siglec-1 were prominent on the cell surface (Figure 4C-4D, see XZ projections below XY). Data with 7-KC treatment thus supports the hypothesis that the CLIC/GEEC pathway is required for endocytosis of VLPs to form the VCC.

**Figure 4.**
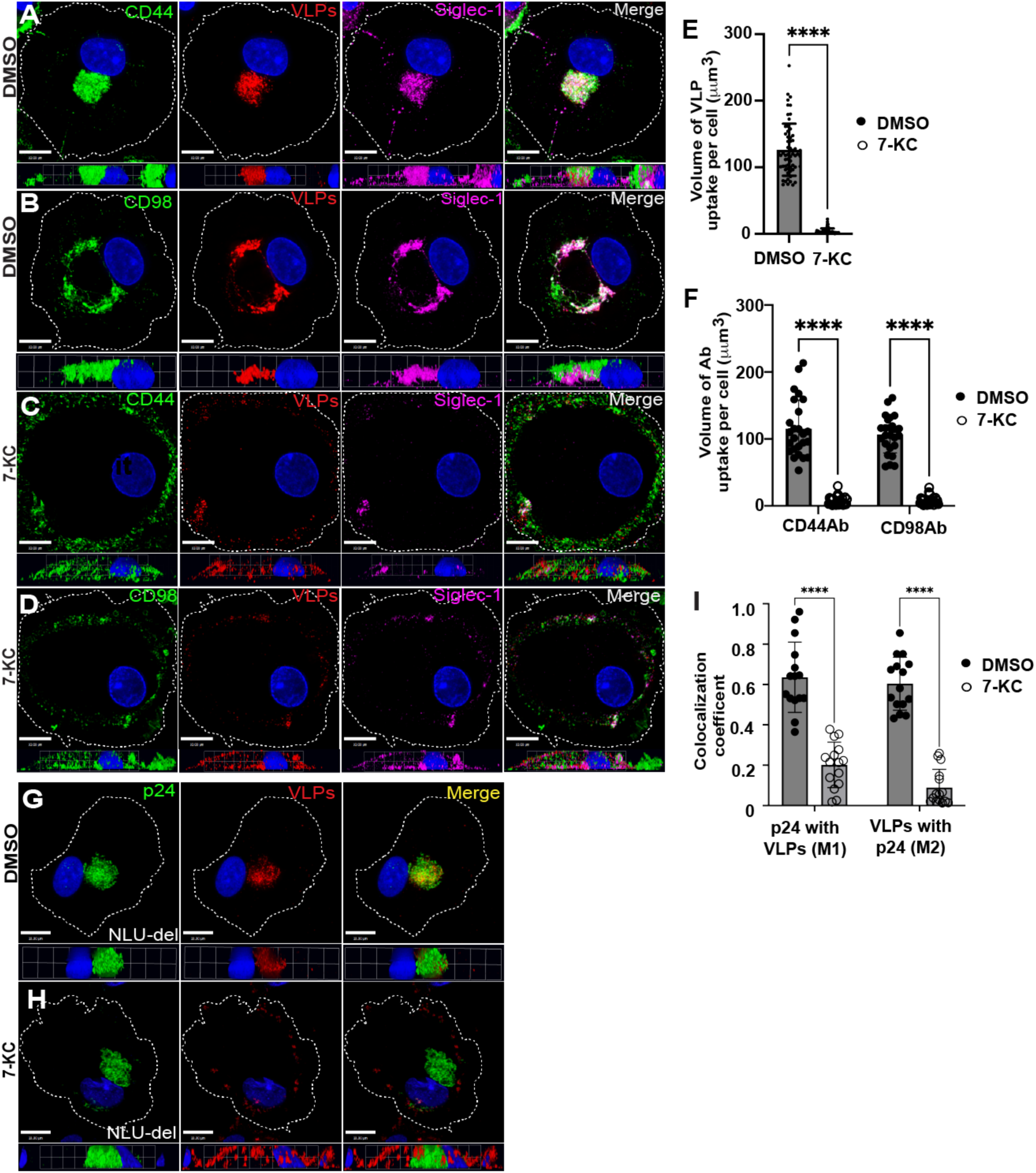
7-KC treatment prevents VLP endocytosis to the VCC. **(A and B)** DMSO control MDMs incubated for 12 hours, followed by addition of mcherry HIV-1 VLPs for 14 hours to allow VLP uptake. Pulse-chase with CD44 Ab (green), Siglec-1 Ab (magenta), and CD98 Ab (green) for 30 minutes was performed prior to fixation of MDMs. Scale bars = 10μm. **(C and D)** MDMs were treated with 7-KC for 12 hours, followed by VLP addition and pulse-chase with antibodies against CD44, CD98, and Siglec-1. XZ projections shown below XY show VLPs are captured and arrested on cell surface. (**E**) Volume of VLP per cell (μm^3^) of control vs 7-KC treated MDMs, mean values ± SD (****P value of <0.0001) from 60 cells/group. **(F)** Volume of CD44 and CD98 uptake per cell for control and 7-KC-treated cells (mean ± SD, ****P value of <0.0001). **(G)** MDMs were infected with NL_Udel_, followed by addition of mcherry VLPs on day 10 post-infection to evaluate VLP (red) internalization to the VCC (green, p24). Note that VLPs reach the pre-formed VCC where they colocalize with mature virions. Scale bar = 10μm. **(H)** MDMs were infected by NL_Udel_ as in (G), then treated with 7-KC on day 10 post-infection at the time of mcherry-VLP addition. Note lack of colocalization of VLP signal with pre-formed VCC (green). (**I**) Colocalization coefficient (M1) of p24 with mcherry VLPs and (M2) of mcherry VLPs with p24 in control vs. 7-KC-treated cells. Values represent measurements from 15 cells/group. ****p <0.001 and ns, not significant (student’s t test).

To further evaluate the involvement CLIC/GEEC pathway in the formation of the VCC, we examined the effects of 7-KC treatment on uptake of VLPs into a pre-formed VCC in HIV-1-infected MDMs. MDMs were first infected with VSV-G-pseudotyped *vpu*-deficient NLUdel, allowing the formation of enlarged VCCs. On day 10 post-infection, MDMs were treated with 7-KC, followed by VLP addition to evaluate endocytosis to the preformed VCC. Note that we used KC-57 antibody to detect mature p24 produced following infection (green), allowing us to distinguish signal from the uncleaved Gag core of mcherry-VLPs (red), as had been previously established (30). Control MDMs demonstrated robust VLP uptake and were concentrated within the VCC of infected MDMs (Figure 4G, quantitation in 4I). In contrast, 7-KC-treated MDMs failed to endocytose VLPs into the pre-formed VCC (Figure 4H, quantitation in 4I). Notably, as had been seen previously with 7-KC treatment followed by VLPs addition, the added VLPs were captured by Siglec-1 but remained predominantly on the surface of the MDMs (Figure 4H, compare to 4G). Thus, experiments using 7-KC indicate that this inhibitor of CLIC/GEEC endocytosis prevents endocytosis of Siglec-1-captured particles to the VCC.

To address the potential that off-target effects of 7-KC could be responsible for the observed inhibition of particle uptake, we confirmed that while 7-KC prevented internalization of CD44 and CD98, it did not inhibit CME, as shown by preserved transferrin uptake in the presence of inhibitor (Supplemental Figure 2A-2D, with quantitation in 2E-2G). Furthermore, 7-KC treatment of MDMs did not inhibit phagocytosis of IgG-opsonized latex beads (Supplemental Figure 2H-2I). Lastly, we observed no significant change in viability of MDMs after 12 hour exposure to 30 μM 7-KC (Supplemental Figure 3M). We conclude that 7-KC did not interfere with endocytosis via the CME pathway or with phagocytosis, and did not diminish cell viability in the time frame utilized in our experiments.

### Inhibition of CLIC/GEEC-mediated endocytosis by IRSp53 depletion prevents particle uptake and VCC formation

Insulin-responsive protein of mass 53kDa (IRSp53) is a BAR domain protein known to interact with CDC42 and ARF1, mediating membrane curvature and interacting with actin regulatory proteins necessary for CLIC/GEEC endocytosis (44, 45). We next examined depletion of IRSp53 as a second method of examining the role of the CLIC/GEEC pathway in VCC formation. IRSp53 was successfully depleted as shown by immunostaining (Figure 5B, G, H) and by western blot (Figure 5C). Control siRNA-treated MDMs demonstrated robust uptake of VLPs, forming the VCC as expected (Figure 5A). Depletion of IRSp53 resulted in marked reduction in the volume of internalized VLPs, with only a few scattered VLPs visible within the cell (Figure 5B, quantitation in 5D). These findings support a model in which inhibition of the CLIC/GEEC pathway inhibits endocytosis of Siglec-1-captured VLPs and eliminates VCC formation.

**Figure 5.**
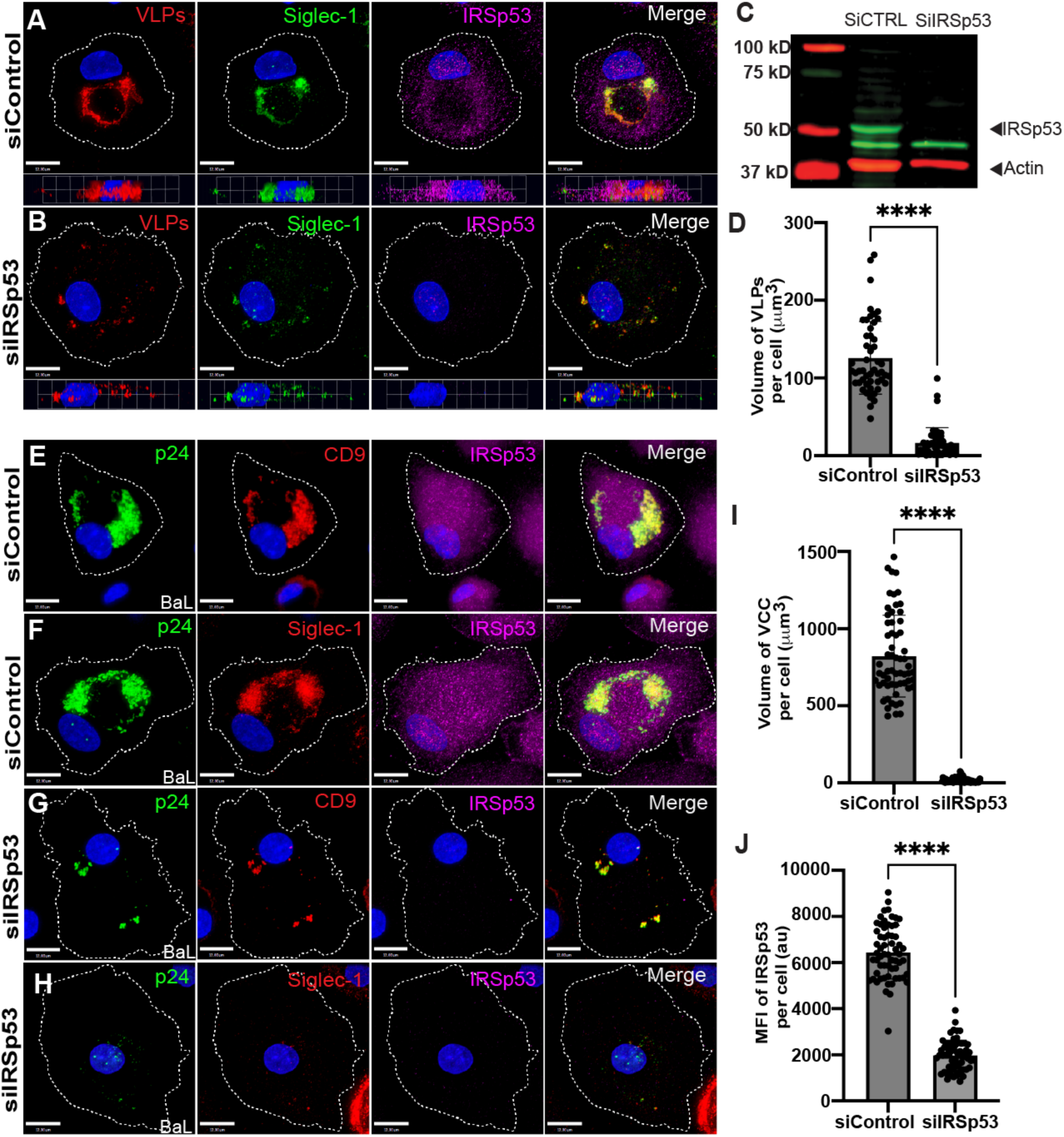
IRSp53 depletion inhibits HIV-1 particle uptake and VCC formation. **(A)** VLP uptake and VCC formation in MDMs transfected with 50nM of control siRNA . XZ profiles are shown below the XY panels. **(B)** VLP uptake in MDMs depleted of IRSp53 using siRNA. Scale bar = 12 μm. **(C)** Representative knockdown of IRSp53 in MDMs that were transfected with either siControl or siIRSp53, as shown by western blot. (**D**) Mean ± SD volume of VLPs per cell, data from a total of 50 MDMs per group. (**E and F**) BaL infected MDMs were transfected with 50nM control siRNA one day following infection, then stained for the indicated markers on day 12. Scale bar = 12 μm. **(G and H)** BaL-infected MDMs transfected with siIRSp53 and stained for the indicated markers on day 12 post-infection. (**I**) Mean ± SD volume of VCC per cell; data from a total of 61 BaL-infected MDMs in each group. (**J**) Mean ± SD measurements of the MFI of IRSp53 expression. Data are representative of of least three independent experiments ****p <0.001 and ns, not significant (student’s t test).

We further examined the role of IRSp53 on VCC formation within HIV-1_BAL_ -infected MDMs by depleting IRSp53 beginning one day post-infection, incubating cells for 12 days, and staining for p24 and VCC markers. Control siRNA-treated MDMs formed large deep VCCs in the MDMs as measured by p24, Siglec-1, and CD9 staining (Figure 5E and 5F). IRSp53-specific siRNA treatment of MDMs resulted in a reduction of IRSP53 staining of >70% as measured by fluorescence intensity (Figure 5J). IRSp53 depletion resulted in a significant reduction in VCC volume within HIV-1 infected MDMs (Figure 5G and 5H), with mean VCC volume reduction from 811 ± 34 mm^!^ to 17± 2 mm^!^ (Figure 5I). We conclude that both chemical inhibition using 7-KC and genetic inhibition through depletion of IRSp53 inhibits endocytosis of HIV-1 particles following capture by Siglec-1, suggesting a role for the CLIC/GEEC pathway in VCC formation within HIV-1-infected MDMs.

### Siglec-1-captured VLPs are endocytosed within membrane tubules together with CLIC/GEEC cargo

Membranous tubules have been shown to emanate from the VCC of infected macrophages (17, 46). The presence of HIV-1 particles within 150–200nm membranous tubules connecting to the PM has been documented by ion abrasion scanning electron microscopy (24). When examining uninfected MDMs, we observed CLIC/GEEC tubules (as indicated by CD98 staining) that extended from the cell surface toward the center of the cell (Figure 6A). Siglec-1 was found primarily in small puncta throughout the cell and on the cell surface, with some limited colocalization with CD98-marked tubules in these uninfected cells (Figure 6A). We hypothesized that the virion-containing tubules previously identified connecting the PM to the VCC may represent Siglec-1-captured virus from the PM within tubules of the CLIC/GEEC pathway. To evaluate this possibility, we performed timecourse experiments in which we pulse-labeled CD98 to mark CLIC/GEEC tubules and then monitored the movement of Siglec-1-captured particles and CD98 from the surface of MDMs into the cell. Remarkably, we found the Siglec-1 captured VLPs concentrated along the CD98+ tubules at early timepoints following VLP addition (Figure 6C, 30 minutes). By 2 hours, the pulsed CD98 marker, together with Siglec-1 and VLPs, formed a more centralized coalescence of membranes in the perinuclear region, forming a VCC-like structure (Figure 6D). By 6 hours post-VLP addition, further coalescence of VLPs, CD98, and Siglec-1 had occurred, and a characteristic horseshoe-like VCC formed (Figure 6E).

**Figure 6.**
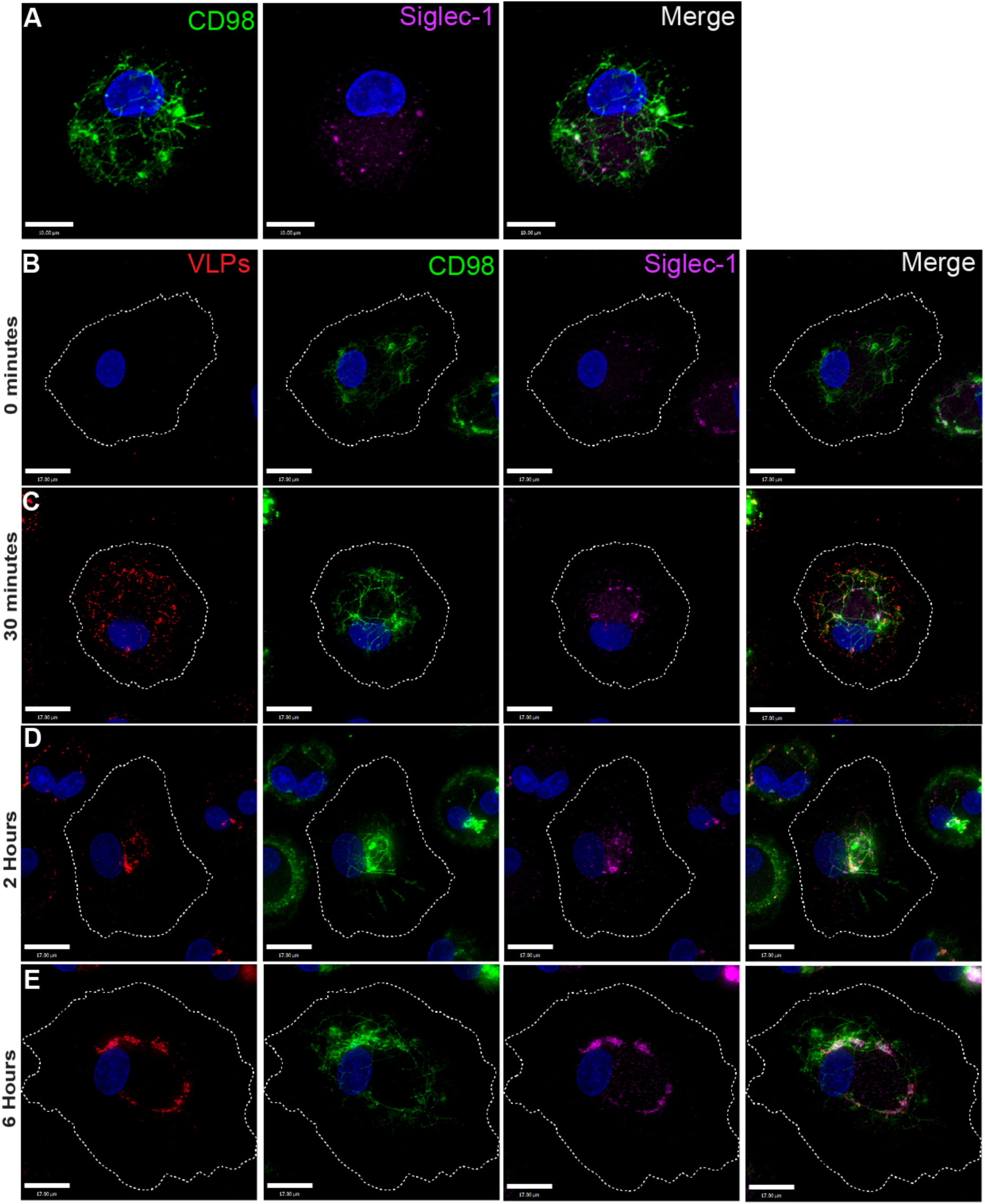
Siglec-1-captured VLPs associate with CLIC/GEEC tubules and move centripetally to form a VCC-like structure. **(A)** MDMs were labeled with CD98Ab (green) for 1.5 hours before fixation and with Siglec-1Ab (magenta). Scale bar = 10 μm. (**B**-**E**) MDMs were pulsed with CD98Ab (green) for 1.5 hours, then mcherry-VLPs (red) were added to the media followed by incubation for 30 minutes (B), 2 hours (D), and 6 hours (E). At the indicated times, MDMs were washed, fixed in warm 4% PFA and secondary antibodies applied for immunostaining of Siglec-1 (purple) and CD98 (green). Scale bar = 17 μm.

### Inhibition of the CLIC/GEEC pathway using 7-KC disrupts tubule formation and prevents internalization of HIV VLPs

The centripetal movement of CLIC/GEEC cargo CD98 together with VLPs shown above suggests that formation of CLIC tubules is an important step in VCC formation. We repeated the previous experiment demonstrating that 7-KC inhibits VCC formation (Figure 4), but utilized pulse-labeling of CD98 followed by a 6-hour chase to monitor tubule movement and VCC formation in the presence of this inhibitor. Figure 7A shows control MDMs with VLPs, Siglec-1, and CD98 tubules coalescing in the center of the cell as previously demonstrated. In contrast, 7-KC treatment led to complete disruption of the tubular architecture marked by CD98 staining (Figure 7B, CD98). VLPs were captured by Siglec-1 under this condition, but remained on the surface of the cell (Figure 7B, VLP and Siglec-1 panels). Together, these findings indicate that the formation of CLIC tubules is an essential step in endocytosis of Siglec-1-captured HIV particles and subsequent VCC formation.

**Figure 7.**
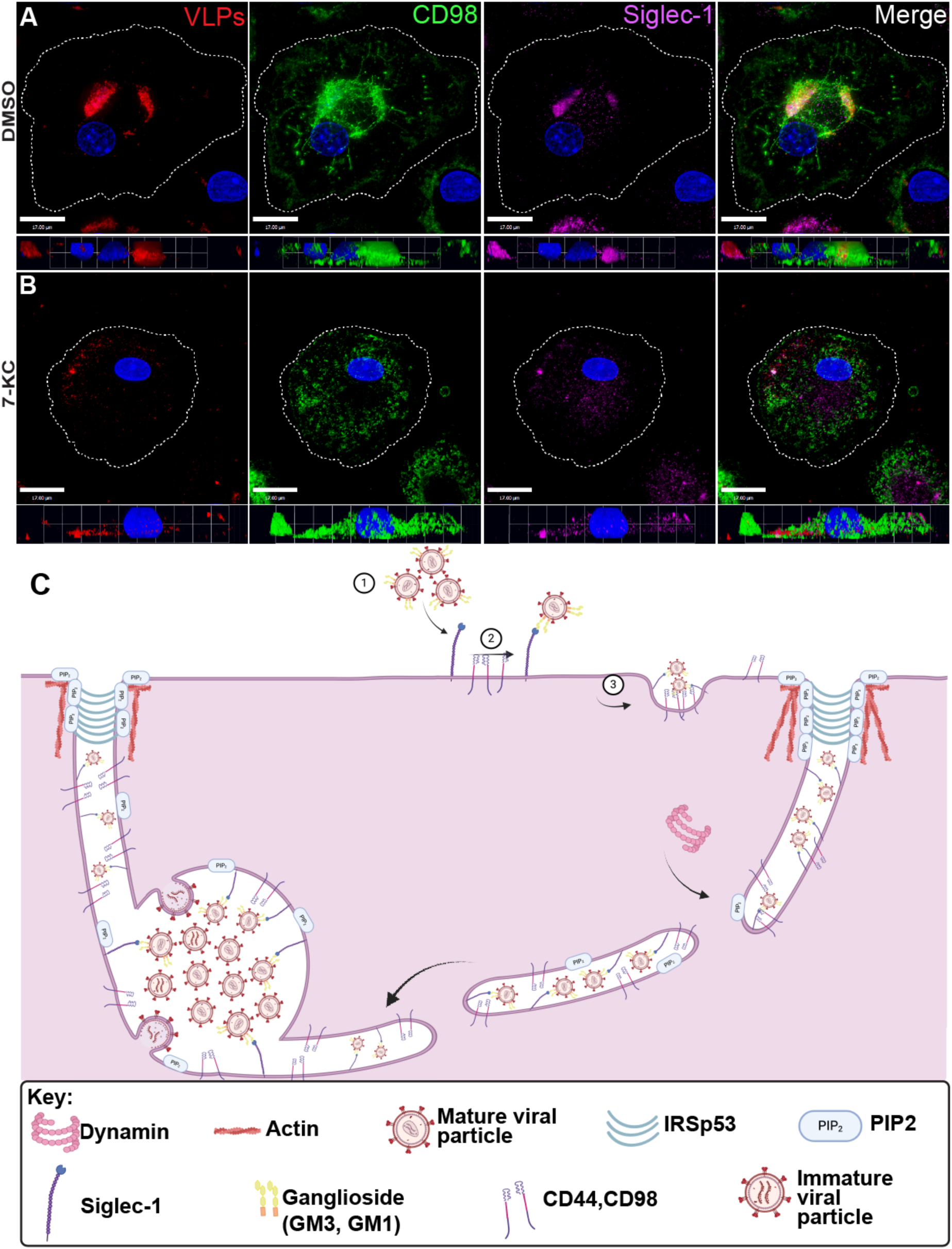
Siglec-1-captured HIV-1 employs CLIC tubules and CLIC/GEEC-mediated endocytosis to form the VCC in Macrophages. **(A)** Control MDMs were treated with DMSO followed by addition of mcherry VLPs. After a 6-hour incubation, CD98 Ab was added for 1.5 hrs, and cells were fixed and stained.**(B)** MDMs were treated with 7-KC followed by VLP addition and incubation as in (A). CD98 Ab was added as for 1.5 hrs prior to fixation. **(C)** Model proposed for the role of CLIC/GEEC in endocytosis of VLPs, and in formation of VCC in infected MDMs. Siglec-1-captured HIV-1 utilizes CLIC/GEEC tubules to enter cells and form the VCC. The role of dynamin-2 remains uncertain but may assist with membrane turnover required during VCC formation. Plasma membrane components diverted to an intracellular site (the VCC) then form an intracellular assembly platform that contributes to expansion of the VCC.

## Discussion

Siglec-1 on the surface of macrophages or dendritic cells (DCs) captures infectious HIV-1 virions or HIV-1 VLPs by attaching to gangliosides on the lipid envelope of the virus, resulting in internalization of particles into the MDMs to form the VCC (28–31). The compartment formed following capture of virions by Siglec-1 can facilitate trans-infection of T cells (28, 29). Siglec-1 nanoclustering occurs in activated DCs, facilitating particle capture and leading to actin rearrangements that promote VCC formation, and this model likely applies to macrophages as well (31). However, the exact mechanisms underlying viral particle endocytosis following capture by Siglec-1 remain unknown. Here we investigated endocytic pathways involved in the internalization of HIV-1 particles for VCC formation in macrophages. We found that following Siglec-1-mediated capture, uptake of particles occurs via the CLIC/GEEC pathway. In contrast to classical CLIC/GEEC endocytosis, however, particle uptake into the VCC also requires dynamin-2 (Figure 7C).

We found strong evidence for the involvement of the CLIC/GEEC pathway in HIV-1 particle uptake and VCC formation. It has been previously described that CD44, a classical marker for the CLIC/GEEC pathway, and is enriched in the VCC (23, 44, 47). We show that a second classical marker of the CLIC/GEEC pathway, CD98, is also highly concentrated in the VCC of the HIV-1 infected MDMs. Chemical inhibition or depletion of essential components of the CLIC/GEEC pathway diminished particle uptake and drastically reduced VCC volume, supporting a central role for the CLIC/GEEC pathway in VCC formation. Remarkably, Siglec-1-captured virions moved together with CD98-marked tubules in a centripetal fashion to form a VCC. We note that other viruses, including adeno-associated virus 2 (AAV2) and Simian Virus 40 (SV40), utilize CLIC/GEEC endocytosis to enter cells (48, 49). Endocytosis of cargo through CLIC/GEEC is characterized by the formation of plasma membrane invaginations that form extended tubules, reminiscent of the tubules previously observed connecting to the VCC of HIV-infected MDMs (47, 50). The tubules demonstrated here are very likely the same tubules previously seen by ion abrasion scanning EM in infected macrophages (24). We suggest from this work that virion-containing tubules seen in these static images in infected macrophages represent Siglec-1-captured viruses that are moving centripetally from the PM toward the VCC, rather than virions leaving the VCC to exit the cell.

The requirement for both dynamin-2 and CLIC/GEEC for endocytosis of HIV-1 particles and formation of the VCC adds some complexity to this pathway, as CLIC/GEEC cargo are typically dynamin-independent. A parallel may be found in Cholera toxin subunit B (CTxB), which also relies on both the CLIC/GEEC pathway and dynamin for entry (51). Although not required for CLIC formation, dynamin has been shown to be recruited to CLICs following plasma membrane scission (41). Recruitment of dynamin to CLICs as they mature may explain the involvement of the BAR domain protein GRAF1, which is known to be a component of tubular endosomes involved in the uptake of GPI-linked proteins and glycosphingolipids (52). GRAF1 binds to dynamin and plays a role in processing of CLIC/GEEC endocytic membranes, suggesting that dynamin itself may facilitate endocytosis of CLIC/GEEC cargo at a step following initial internalization from the plasma membrane (53) . We propose a model in which Siglec-1-mediated capture of HIV-1 particles and nanoclustering of Siglec-enriched membranes leads to tubulation of the membrane inwards to form the CLIC. After the CLIC has matured into the GEEC, dynamin-2 may be required for a membrane scission step required for the formation of the VCC (depicted in Figure 7C). Diversion of plasma membrane components including PI(4,5)P_2_ to the VCC through this pathway may then set the stage for subsequent HIV assembly events to occur on the VCC membranes. In support of this idea, CLIC/GEEC endocytosis occurs in regions of the plasma membrane that are enriched in PI(4,5)P_2_ or PI(3,4,5)P_3_, and the CLIC/GEEC regulator GRAF1 also binds to PI(4,5)P_2_-enriched membranes (54). VCC membranes are enriched in PI(4,5)P_2_, thus creating “assembly platforms” within the cell that have combined with late endosomal components to form the VCC (55). The model outlined here proposes that the PI(4,5)P_2_-enriched membranes of the VCC are derived from the PM through the CLIC/GEEC pathway. This model provides an origin for the tubules connecting VCCs to the plasma membrane, and a connection between initial Siglec-1-mediated particle capture and delivery to the VCC. The prominent presence of CLIC/GEEC cargo (CD44, CD98) in the VCC, the movement of tubular membranes together with HIV VLPs to form a VCC, and the inhibition of VCC formation by inhibition of CLIC/GEEC endocytosis all support this model.

The VCC is a compartment known to facilitate trans-infection of lymphocytes, and therefore contribute to HIV spread and pathogenesis. This compartment may also serve as a reservoir of infectious virions within tissue macrophages and contribute to HIV persistence in the presence of antiretroviral therapy. Identification of the important role the CLIC/GEEC pathway plays in VCC formation will facilitate future efforts to eradicate infectious virus from this compartment.

## Materials and methods

### Ethics statement

Human blood for the preparation of monocyte-derived macrophages and other experiments in this work was obtained from volunteer donors and was de-identified prior to handling by the investigators. Informed consent was obtained from participants. Blood was collected under a protocol approved by the CCHMC Institutional Review Board.

### Isolation and maturation of MDMs

Human peripheral blood mononuclear cells (PBMCs) were isolated from human blood using Ficoll-Hypaque gradient centrifugation. PBMCs from the buffy coats were pooled and extensively washed with PBS. Monocytes were enriched by negative selection method using Miltenyi Pan monocyte isolation kit (Miltenyi Biotec Inc). Enriched monocytes were plated on either multiwell chambered coverglass (Cellvis) or poly-D-lysine coated 35mm^3^ Mattek dishes (MatTek Corporation) or poly-D-lysine coated 6 well plates (Corning) and cultured in RPMI 1640 media supplemented with 10% FBS, 100 U/ml penicillin, 100 ug/ml streptomycin and 2mM Glutamine. The cells were matured in the presence of 5 ng/ml GM-CSF for 7 days to facilitate maturation into monocyte derived macrophages (MDMs). Following maturation, the MDMs were stimulated with 5 ng/ml GM-CSF (PeproTech Cat no 300-03) and with 2000 U/ml of universal Type1 Interferon (PBL Assay Science Cat. No. 11200-2).

### Reagents, inhibitors and antibodies employed

Dyngo4a (Cat no AB120689) was obtained from Abcam and used at a concentration of 20 μM. Nps2143 (Cat no SML0362) was obtained from Sigma-Aldrich and used at a concentration of 5 μM. 5-(N-Ethyl-N-isopropyl)amiloride (Cat no. A3085-25MG) was obtained from Millipore Sigma and was used at a concentration of 25 μM. Pitstop 2.0 (Cat no SML1169) was obtained from Sigma-Aldrich and Pitstop negative control (Cat no AB120688) was obtained from Abcam and used at a concentration of 5 μM. 7-Keto Cholesterol (7-KC) (Cat. No. 16339) was obtained from Cayman Chemical Company and was used at 30 μM. Fluorescently labeled Alexa Fluor 647 and Alexa Fluor 488 human transferrin were purchased from Jackson ImmunoResearch (Cat. No. 009-600-050 and 009-540-050) and employed at a dilution of 1:100. Fluorescently labeled 70 kDa Dextran (Cat. No. D1822) was purchased from Thermo Fisher Scientific and was used at a concentration of 100 μg/ml. Mouse monoclonal antibodies to human CD9 (Cat. No.312102) diluted to 1:250, human Siglec-1 (Cat. No 346002) diluted to 1:200 and Alexa Flour 647 anti-human CD169 (sialoadhesin, Siglec-1) (Cat No. 346005) diluted to 1:150 were from Biolegend. FITC-KC57 was obtained from Beckman Coulter (Cat. No. 66046650) and was used at a dilution of 1:500. EPS15 antibody (Cat. No. NBP1-89221) purchased from Novus Biologicals was used for immunoblotting. Rabbit Polyclonal anti-human FCHO2 antibody (Cat. No. PA5-31696) from Thermo Fisher Scientific was used for immunoblotting (dilution 1:1,000). Mouse monoclonal antibody to human CD44 (Cat. No. ab6124) from Abcam was used in immunofluorescence studies at 1:500 dilution. Anti-human CD98 antibody (Cat. No. 315602, 1:250) and FITC Anti-human CD98 Antibody (Cat No. 315603, 1:150) were from Biolegend. Rabbit polyclonal Anti-IRSp53 antibody (Cat. No. ab126057) from Abcam was used for immunoblotting (1:5000) and immunofluorescence (1:500). BV421 Mouse Anti-Human CD98 antibody (Cat. No. 744502) from BD bioscience was used at a 1:100 dilution for pulse-chase experiments. Mouse (Cat. No. Ma5-11869) and Rabbit (Cat. No. A20266-100UL) actin antibodies were purchased from Thermo Fisher Scientific and Millipore Sigma for western blotting (1:5000). Secondary antibodies included Alexa Fluor 488-conjugated goat anti-mouse, Alexa Fluor 488-conjugated goat anti-Rabbit, Alexa Fluor 647-conjugated goat anti-mouse and Alexa Fluor 647-conjugated goat anti-Rabbit (Invitrogen) at a dilution of 1:1,000.

### VLP production

HIV-1 mcherry VLPs were generated by transient transfection of HEK 293T cells with a pVRC-3900 and pVRC/ GAGOPT-mcherry at a ratio of 4:1 as previously described (30). The HIV-1 Pr55^Gag^ construct, pVRC-3900, is an expression plasmid encoding a codon-optimized HIV-1 Gag polyprotein and was kindly provided by Gary Nabel (VRC, NIH). HEK293T cells were transfected using JET prime reagent (PolyPlus). VLPs were harvested 48 hours after transfection, supernatants clarified and concentrated by centrifugation through a 20% sucrose cushion. VLPs were resuspended in ice cold serum free RPMI 1640 and the aliquots were stored at -80°. For use, the VLP aliquot was thawed and filtered through a 0.45um syringe filter before addition to MDMs.

### Viral stock generation and MDM infection

pNL_Udel_ proviral plasmid was obtained from Klaus Strebel, NIAID, NIH. VSV-G-pseudotyped HIV-1 NLUdel stocks were generated by transfection of 293T cells using Jetprime transfection reagent (Polyplus). 36 hours after transfection, viral supernatant was harvested, filtered through a 0.45 μm filter and stored at -80 °C. The viral titer was determined using a TCID_50_ assay performed in TZM-bl indicator cells (obtained through the NIH HIV Reagent Program, Division of AIDS, NIAID, NIH: TZM-bl Cells, ARP-8129, contributed by Dr. John C. Kappes, Dr. Xiaoyun Wu and Tranzyme Inc). Primary HIV-1 isolate BaL stocks were prepared as follows: Human peripheral blood mononuclear cells (PBMCs) were resuspended in RPMI 1640 supplemented with 20% heat-inactivated fetal bovine serum and 50 μg/ml gentamicin (RPMI 1640-GM). Primary HIV-1 isolates were propagated in PBMCs stimulated with 5 μg/ml phytohemagglutinin (PHA) and 5% interleukin 2 (IL-2). The IL-2/ PHA-stimulated cells were infected using a high-titer seed stock of virus minimally passaged in PBMCs, derived from a viral stock obtained through the NIH HIV Reagent Program (HIV-1 Ba-L, ARP-510, contributed by Dr. Suzanne Gartner, Dr. Mikulas Popovic and Dr. Robert Gallo). 1 ml of virus was transferred to the flask containing freshly stimulated PBMCs and incubated overnight at 37 °C in 5% CO_2_. Cells were washed extensively and resuspended in 30 ml of RPMI-GM with IL-2. The virus-containing supernatants were collected, clarified by centrifugation, and filtered through a 0.45 μm filter and stored at -80 °C.

For titering of viral stocks, TZM-bl cells were incubated for 48 hours after infection with viral supernatant and 100 μl of supernatant was removed from each well prior to the addition of 100 μl of Britelite (Perkin-Elmer) substrate. Measurement of infectivity was performed using 150 μl of cell/substrate mixture in a black 96 well solid plate and measurement of luminescence was performed using a plate luminometer. Human MDMs were infected with the viral stock at a MOI of 3 for NL_Udel_ experiments and a MOI of 0.5 for BaL experiments.

### Immunofluorescence staining and Imaging

MDMs were fixed with 4% PFA in PBS for 10 min at room temperature and permeabilized with 0.1% Triton 100 for 5 minutes, then washed thoroughly with PBS. Fixed cells were blocked with Dako blocking buffer supplemented with 1 μg/ml human IgG. Cells were then incubated with antibodies against Siglec1, CD9, CD44, CD98, IRSp53 or KC57 in Dako antibody diluent (Dako) overnight at 4 °C. Cells were washed thoroughly before incubations with appropriate secondary antibodies if not incubated with a primary labeled antibody. For visualization of the nucleus, the MDMs were stained with DAPI (4^’^-6’-diamidino-2-phenylindole) at 300 nM in PBS. Imaging was performed on a Deltavision RT deconvolution microscope (Applied Precision/ Leica Instruments), and data analysis was performed using Volocity software (Perkin-Elmer/Quorum Technologies). The number of cells quantified for each experimental arm is indicated in the respective figure legends.

### VLP uptake in chemical inhibition experiments

VLP uptake assays were performed in GM-CSF-matured MDMs after stimulation with IFN-U. HIV-1 mcherry VLPs or both HIV-1 mcherry VLPs or HIV-1 Gag-GFP VLPs were added to MDMs in the presence of the indicated inhibitors or controls. For the experiments with Dyngo4a, the MDMs were first serum starved for 30 minutes at 37 °C prior to addition of HIV-1 Gag-GFP VLPs. After serum starvation, 400 ng of HIV-1 Gag-GFP VLPs were added to the MDMs for the next 5 hours. MDMs were washed with serum free RPMI to remove residual HIV-1 Gag-GFP VLPs. Next, the MDMs were treated with either 20μM of Dyngo4a or 20μM DMSO control for 30 minutes at 37 °C. After 30 minutes, 400 ng of HIV-1 mcherry VLPs in serum free RPMI with either 20 μM of Dyngo4a or 20 μM DMSO control were added to the MDMs for 2.5 hours. During the last 30 minutes of the HIV-1 Gag-mcherry VLP uptake, the MDMs were incubated with Alexa Fluor 647-conjugated human transferrin. After the internalization, the MDMs were rinsed with PBS and then fixed with warm 4% PFA in PBS for 10 minutes at 37 °C and were rinsed with 100 nM glycine in PBS and processed for immunofluorescence staining. For experiments containing 7-KC, MDMs were treated with either 30 μM of 7-KC or 30 μM DMSO control for 12 hours. After this treatment, 400 ng of HIV-1 mcherry VLPs in serum-free RPMI with either 30 μM of 7-KC or 30 μM DMSO control were added to the MDMs for 14 hours. During the last 30 minutes of the VLPs uptake, MDMs were incubated with FCR blocker and CD44Ab or with FCR blocker and FITC-CD98Ab at 37 °C. The MDMs were then washed with PBS and fixed, stained for Siglec-1 and imaged as described above.

### CD98 antibody pulse-chase in MDMs

MDMs were plated on either 35 mm^3^ Mattek dishes (MatTek corporation) or multiwell chambered coverglass (Cellvis). On day 6 after maturation, MDMs were treated with 2000 U/ml IFN-U for 2 days. On day 2, the MDMs were serum starved for 30 minutes at 37 °C. MDMs were then next incubated with BV421 anti-CD98 antibody against the endogenous CD98 protein for 1.5 hours at 37 °C to allow for internalization of the antibody bound CD98. After the pulse-chase with CD98 antibody, MDMs were rinsed twice with serum free RPMI medium, followed by addition of 400 ng of HIV-1 mcherry VLPs, and incubated for 30 minutes, 2 hours and 6 hours after the addition of VLPs. To obtained time point 0 after VLPs addition, some MDMs were fixed immediately after VLPs addition with warm 4% PFA in PBS for 10 minutes at 37 °C. After fixation, MDMs were rinsed with 100 nM glycine in PBS. MDMs were then permeabilized with 0.05% Triton 100 for 5 minutes and washed with 0.025% Triton 100 for 5 minutes. MDMs were blocked with Dako blocking buffer supplemented with 1 μg/ml human IgG. MDMs were then incubated with Alexa Flour 647 anti-human CD169 antibody in Dako antibody diluent (Dako) overnight at 4°. After overnight staining, the MDMs were washed with 0.025% Triton multiple times. For visualization of the nucleus, MDMs were stained with DRAQ5 at 1:250 dilution in PBS for 15 minutes at room temperature. For the CD98 antibody pulse-chase experiment with 7-KC inhibitor addition, MDMs were treated with either 30 μM of 7-KC or 30 μM DMSO control for 12 hours. After this, 400 ng of HIV-1 mcherry VLPs with either 30 μM of 7-KC or 30 μM DMSO control were added to the MDMs for the next 6 hours. During the last 1.5 hours of the VLPs uptake, the MDMs were incubated with BV421 anti-CD98 antibody against the endogenous CD98 protein to allow for internalization of CD98. After the CD98 antibody pulse-chase, the MDMs were fixed with warm 4% PFA in PBS for 10 minutes at 37 °C.

### Off-target effects assays in MDMs

MDMs were plated on either 35 mm^3^ Mattek dishes (MatTek corporation) or multiwell chambered coverglass (Cellvis). After maturation, MDMs were treated with 2000 U/ml IFN-U for 2 days. MDMs were pre-incubated with inhibitors 7-KC for 12 hours, Pitstop 2.0 for 30 minutes, Dyngo4a for 30 minutes, or controls such as DMSO or Pitstop Negative control for 30 minutes in serum-free RPMI. After the pre-incubation, MDMs were treated with different markers for 30 min for internalization. CD98 Ab and CD44 Ab uptake was used as a marker for CLIC/GEEC pathway, human labeled transferrin (5 ug/ml) for clathrin-mediated endocytosis and fluorescently labeled 70kDa dextran (100 μg/ml) for macropinocytosis. MDMs were then washed with either PBS or an acidic buffer 0.2M glycine with 0.15M Nacl before fixation with 4% PFA.

### Phagocytosis assay

MDMs were treated with 2000 U/ml IFN-U for 2 days post maturation. On day 2, MDMs were treated with either 30 mM of 7-KC or DMSO for 12 hours. After the pre-incubation, the MDMs would have a change in medium to serum free RPMI that contained either 30 mM of 7-KC or DMSO with HIV-1 mcherry VLPs for 14 hours. After 14 hours, latex beads coated with fluorescently labeled IgG from the phagocytosis assay kit (Cayman Chemicals Cat. No. 500290) were added to the MDMs for 1 hour at 37 °C. Then, the MDMs were washed with assay buffer to remove any beads from the MDM cell surface. After washing, the MDMs were fixed and stained as outlined above.

### Knockdown experiments in MDMs with VLP uptake

SiCtrl (12935300), siDnm2 (4390824 ID: s4212) and siEps15 (4392420 ID: s4775) were obtained from Thermo Fisher Scientific and both siFCHO2 (Cat. Number sc-91916) and siIRSp53 (Cat. No. sc-60863) were obtained from Santa Cruz Biotechnology. For knockdown experiments, MDMs were treated with either 50 nM of siCtrl, siDnm2, siFCHO2, siIRSP53 or siEps15 using Lipofectamine RNAiMax reagent (Life Technologies) using the manufacturer’s protocols. After 7 hours, an equal amount of serum containing medium was add to the transfection medium and on day 1 post-transfection, the medium removed and replaced with macrophage medium. 3 days post-transfection, MDMs treated with either siDnm2 or siEps15 received a second round of siRNA treatment, and on day 5, the si-MDMs were then treated with 2000 U/ml universal Type1 Interferon for 16 hours. On day 6 following the initial siRNA transfection, HIV-1 mcherry VLPs in serum free medium were added, and cells incubated for an additional 14 hours. The MDMs were then fixed and stained for immunofluorescence imaging.

### Knockdown experiments in MDMs with HIV-1 infections

MDMs were treated with VSV-G pseudotyped HIV-1 NL_Udel_ at MOI of 2 or with BaL at an MOI of 0.5 for 16 hours. After HIV-1 infection, MDMs received either 50 nM of SiCtrl, siDnm2, and siEps15 as described above. After 7 hours, an equal amount of serum containing medium was add to the transfection medium and on day 1 post-transfection, the medium removed and replaced with macrophage medium. On day 6 post-transfection, MDMs treated with either siDnm2 or siEps15 received a second round of siRNA treatments as described earlier. MDMs infected with NL_Udel_ were fixed on day 10 post-infection, and MDMs infected with BaL were fixed on day 12 post-infection.

### Western blotting

MDM lysates were prepared by lysing in RIPA buffer containing protease inhibitors for 30 min on ice, followed by clarification by centrifuging at 15000 RPM on a tabletop centrifuge at 4 °C for 30 min. Protein concentration of the lysates was measured using the DC protein assay kit (BioRad 5000111) following the manufacturer’s instructions. 40 μg of total protein was loaded onto precast 4-12% Bis-Tris NuPAGE gels (Thermo Fisher Scientific). Protein was transferred to nitrocellulose membrane using the semi-dry trans Blot transfer (BioRad) following manufacturer’s instructions. The membrane was blocked, probed with appropriate primary and secondary antibodies, and imaged using the LiCOR infrared imaging system.

## Acknowledgments

We thank Matthew Kofron of the CCHMC Bio-imaging and Analysis Facility for helpful input regarding image analysis.

